# Lateral habenula M2 muscarinic receptor control of neuronal activity and cocaine seeking behavior

**DOI:** 10.1101/2021.07.24.453572

**Authors:** Clara I.C. Wolfe, Eun-Kyung Hwang, Agustin Zapata, Alexander F. Hoffman, Carl R. Lupica

**Affiliations:** Computational and Systems Neuroscience Branch, Electrophysiology Research Section, Baltimore, MD 21224; Department of Behavioral Neuroscience, Oregon Health Sciences University, Portland, OR 97239

## Abstract

The lateral habenula (LHb) plays a central role in balancing reward and aversion by opposing the contributions of brain reward nuclei. Using a rat cocaine self-administration model, we previously found that LHb inhibition or non-selective blockade of LHb muscarinic acetylcholine receptors (mAChRs) led to persistent cocaine seeking despite its signaled unavailability. As understanding roles for the LHb and cholinergic signaling in behavioral control is important to psychiatric illness and addiction, we examine how mAChRs act on LHb neurons using *in vitro* electrophysiology. We find that different groups of LHb neurons are depolarized or hyperpolarized by the cholinergic agonist carbachol (CCh), and that CCh could inhibit GABAergic and glutamatergic synaptic inputs to these cells. Presynaptic CCh effects were reversed by the M2 mAChR (M2R) antagonist AFDX-116, but not by pirenzepine, an M1R antagonist. Contemporaneous measurement of CCh effects on synaptic inhibition and excitation in LHb neurons showed a smaller effect on inhibition, suggesting a net shift in synaptic integration toward greater inhibition by mAChRs. Synaptic currents elicited by light-activation of ventral tegmental area (VTA) axons in the LHb, following channelrhodopsin-2 transfection of VTA, were also inhibited by M2Rs, suggesting the VTA as at least one M2R-sensitive LHb afferent. Finally, Go-NoGo cocaine seeking studies showed that blockade of LHb M2Rs, and not M1Rs, triggered continued cocaine seeking. These data identify LHb M2Rs as a potential control point of LHb function that enables withholding responses for cocaine and define cellular mechanisms through which mAChRs modulate LHb activity.

## Introduction

The lateral habenula (LHb) forms part of the habenular complex, a small structure located at the posterior dorsomedial end of the epithalamus (Sutherland, 1982; Kim and Chang, 2005), that is characterized as a hub integrating information from the limbic forebrain before relaying it to major midbrain monoaminergic nuclei (Sutherland, 1982). In a seminal study by Matsumoto & Hikosaka (2007), the LHb was shown to be activated when a received reward was smaller than expected (a negative prediction error), and inhibited during positive reward outcomes (Matsumoto and Hikosaka, 2007), and these changes in LHb neuron activity oppose that of midbrain dopamine (DA) neurons (Matsumoto and Hikosaka, 2007). Later, it was shown that the LHb exerts this control over DA neurons via excitation of the rostromedial tegmentum (RMTg) (Jhou et al., 2009; Balcita-Pedicino et al., 2011; Barrot et al., 2012), a collection of inhibitory neurons forming synapses with DA cells (Christoph et al., 1986), and modulating their influence on motivated behaviors (Stopper et al., 2014).

In this manner the LHb is thought to play a crucial role in processing both rewarding and aversive stimuli by its inverse influence on the brain DA reward system through modulation of the strength and duration of reward signals (Koob and Le Moal, 2008). Not surprisingly, this system has been implicated in substance use disorders (SUDs) and has been more generally characterized as a neural substrate in opponent process theory as it relates to reward and motivation (Solomon, 1980; Ettenberg et al., 1999). Specifically, evidence suggests that many drugs that are initially rewarding also possess aversive properties, and this can promote drug seeking and use through the desire to mitigate or avoid these aversive effects. Thus, in rodents, cocaine is initially rewarding followed by strong aversion and avoidance of drug-paired environments (Ettenberg et al., 1999; Lecca et al., 2017), and biphasic LHb neuron activity that parallels the rewarding and aversive phases have been described (Matsumoto and Hikosaka, 2007; Jhou et al., 2013; Neumann et al., 2014). In addition to these studies, LHb electrical stimulation influences reward-based decision making (Stopper et al., 2014), and inhibition of LHb activity can promote impulsive cocaine seeking (Zapata et al., 2017) and disrupt choice bias in rats (Stopper and Floresco, 2014). Thus, based on its ability to regulate motivational valence, the LHb has been proposed to play a critical role in both SUDs and major depressive disorder (MDD) (Shabel et al., 2014; Yang et al., 2018).

As there is limited evidence for the presence of GABAergic interneurons within the LHb and most of the neurons within this structure are glutamatergic (Brinschwitz et al., 2010; Aizawa et al., 2012), inhibitory control of the LHb is largely derived from extrinsic sources (Zhang et al., 2016; Wagner et al., 2017). Inhibitory LHb afferents from the lateral preoptic area (Barker et al., 2017), lateral hypothalamus (Lecca et al., 2017; Lazaridis et al., 2019), ventral pallidum (Faget et al., 2018), ventral tegmental area (VTA)(Stamatakis et al., 2013; Root et al., 2014a), and medial dorsal thalamus (Webster et al., 2020) have all been recently characterized. When activated, these afferents suppress LHb output and are generally rewarding in behavioral assays. In contrast, excitatory inputs to the LHb from the lateral preoptic area (Barker et al., 2017), entopeduncular nucleus (EPN) (Li et al., 2021) or ventral pallidum (Faget et al., 2018) increase excitatory LHb output and are aversive in rodents. It is also notable that some LHb inputs, such as those from VTA and EPN, can co-release GABA and glutamate (Root et al., 2014b; Shabel et al., 2014; Root et al., 2018).

Understanding the diversity of these afferents and their integration at the neuronal level is important for understanding ways in which the LHb participates in behavioral control. In particular, reduced GABAergic inhibition of the LHb favors excitation and is described in a rodent model of depression (Shabel et al., 2014), during drug withdrawal (Meye et al., 2016), and is associated with increased attributions of negative reward valence and enhanced stress sensitivity (Shabel et al., 2014; Meye et al., 2016; Root et al., 2018). Similarly, prolonged excitability of the LHb during cocaine withdrawal has been associated with an imbalance in GABAergic to glutamatergic transmission from EPN inputs (Neumann et al., 2014; Meye et al., 2016).

Although glutamatergic and GABAergic afferents are important regulators of LHb activity, other neurotransmitters and modulators likely also play a central role. Thus, there is evidence for robust cholinergic innervation of the LHb (Woolf and Butcher, 1986; Geisler et al., 2003; Wagner et al., 2016), with the vesicular acetylcholine transporter (vAChT), acetylcholinesterase (AChE), and mAChR mRNA all highly represented in the medial LHb (mLHb) (Wagner et al., 2016). Moreover, endogenous cholinergic signaling in the LHb appears to be relevant to behavioral control as we have shown that LHb mAChRs are necessary to permit response inhibition for cocaine in a Go/NoGo discrimination task in rats (Zapata et al., 2017). However, despite this apparent cholinergic influence on behavioral control, the mechanisms through which mAChRs influence LHb neuron activity are poorly understood. Here, using *in vitro* electrophysiological techniques we examine the mechanisms in which mAChRs control mLHb neuron activity and use a model of impulse control in rats to evaluate how this influences cocaine motivated behaviors.

## Materials and methods

### Subjects

Male and Female Long Evans rats (Charles River Laboratories, Wilmington, MA) aged 4-6 weeks were housed 2–4 same sex animals per cage in an Association for Assessment and Accreditation of Laboratory Animal Care (AAALAC) International accredited facility. They were maintained in a temperature- and humidity-controlled environment with *ad libitum* food and water. Rats used in electrophysiological experiments were housed under standard lighting conditions (lights on 6:00 am-6:00 pm). For behavioral experiments, rats were housed under a reverse 12 h light/dark cycle, with experiments performed during the dark cycle. All experimental procedures were approved by the Institutional Animal Care and Use Committee (ACUC) of the National Institute on Drug Abuse Intramural Research Program, National Institutes of Health (Rockville, MD) and conducted in accordance with PHS policy, NIH Guidelines, and the Guide for the Care and Use of Laboratory Animals (National Research Council (U.S.). Committee for the Update of the Guide for the Care and Use of Laboratory Animals. et al., 2011).

### Electrophysiology

Animals were anesthetized with isoflurane and decapitated using a guillotine. The brains were extracted and transferred to ice-cold HEPES-modified cutting solution (in mM: NaCl, 92; KCl, 3; NaH_2_PO_4_,1.2; NaHCO_3_, 30; HEPES, 20; Glucose, 25; Ascorbic acid, 5;; MgCl_2_, 10; CaCl_2_, 0.5). The tissue was glued to the stage of a vibrating tissue slicer (Leica VT1200S, Leica Biosystems, Wetzler, Germany). Coronal slices (280 μm) containing the LHb, corresponding to approximately 3.3 mm to 4 mm posterior to bregma (Paxinos and Watson, 2007) were transferred to a holding chamber containing normal aCSF consisting of (mM): NaCl,126; KCl, 3.0; MgCl_2_, 1.5; CaCl_2_, 2.4; NaH_2_PO_4_, 1.2; glucose, 11.0; NaHCO_3_, 26, saturated with 95% O2/5% CO2, at 35 °C for 15-20 min, then maintained at room temperature. A hemisected brain slice was submerged in a low-volume recording chamber (∼170 μl; Warner Instruments, Holliston MA) integrated into a fixed stage of an upright microscope (Olympus BX51WI, Waltham MA) and continuously perfused with warm (30–33 °C) aCSF at 2 ml/min using a peristaltic pump. The aCSF was warmed using an inline solution heater (TC-324B, Warner Instruments). Drugs were prepared as stock solutions in H_2_O or DMSO and diluted in aCSF to the indicated concentrations. Visualization of LHb neurons was performed using differential interference contrast microscopy with infrared illumination. Recordings were performed in a region corresponding primarily to the parvocellular subnucleus of the LHb (LHbMPc) or its central subnucleus (LHbMC) of the medial division (Geisler et al., 2003). Whole-cell voltage clamp recordings were performed using a MultiClamp 700B (Molecular Devices, San Jose CA), WinLTP software (WinLTP Ltd, Bristol, UK), and an A/D board (PCI-6251, National Instruments, Austin TX). Series resistance was monitored throughout recordings using brief hyperpolarizing steps (−10 mV, 200 ms), and cells demonstrating >20% change in access were excluded from analyses.

For recording spontaneous IPSCs, recording electrodes (4–6 MΩ) were filled with (in mM): KCl, 145; HEPES, 10; EGTA, 0.2; MgCl_2_, 2; Mg-ATP, 4; Na_2_-GTP, 0.3; Na_2_-phosphocreatine, 10; pH 7.2 with KOH. For spontaneous EPSCs and electrically evoked or light-evoked IPSCs, recording electrodes were filled with (in mM): K-gluconate, 140; KCl, 5; HEPES, 10; EGTA, 0.2; MgCl_2_, 2; Mg-ATP, 4; Na_2_-GTP, 0.3; Na_2_-phosphocreatine, 10; pH 7.2 with KOH. For neurons clamped at 0 mV (below), QX-314 (1 mM) was added to the intracellular solution. Except when both EPSCs and IPSCs were recorded in the same neuron, IPSCs were pharmacologically isolated using DNQX (10 µM), and EPSCs were isolated in the presence of picrotoxin (100 µM). In some experiments, gabazine (10 µM) or DNQX (10 µM) was bath applied at the end of recordings to confirm the nature of the synaptic response.

Electrically evoked responses were obtained by positioning the tips of a bipolar stimulating electrode (FHC, Bowdoin, ME) on the surface of the brain slice 150-200 µm from the recording electrode within the LHb. Stimulation intensity was adjusted to elicit a response 30-50% of the peak amplitude. For light-evoked currents, a 473 nm DPSS laser (OEM Laser BL-473-00200, Midvale UT) was used to deliver a single light pulse (5 ms), collimated through a 40x objective using a fiber optic adaptor (IS-OGP; Siskiyou, Grants Pass, OR). Light-evoked responses were obtained every 30s, and spontaneous synaptic events were collected as continuous recordings before and during drug application. Continuous event recordings were analyzed offline in 2-minute epochs (before and during drug application). Event detection of spontaneous synaptic currents was performed using WinEDR (University of Strathclyde, Glasgow, UK), using a template-based matching algorithm (Clements and Bekkers, 1997). Typical detection parameters for EPSCs utilized rise times of 0.3 ms and decay times of 3 ms, and individual events were confirmed by visual inspection. Excitatory-Inhibitory (E-I) ratios were calculated by measuring electrically-evoked EPSCs at -70 mV (inward currents) and IPSCs at 0 mV (outward currents) within the same cell, with the stimulation intensity adjusted to produce an EPSC that was approximately 50% of maximum. The E-I ratio was defined as the proportion of excitatory synaptic current to the total synaptic current recorded in each cell, or E/E+I, using the area under the curve (AUC) of 6-10 events averaged during the control pre-drug period and during drug application (Antoine et al., 2019).

### Surgery

#### Virus infusions

Rats were positioned in a stereotaxic frame (Kopf Instruments, Tujunga CA) and initially anesthetized with isoflurane (4% delivered at a flow rate of ∼1 L/min O_2_) using a calibrated vaporizer. Thereafter, anesthesia was maintained with 1-1.5% isoflurane delivered at a flow rate of ∼0.2 L/min O_2_. Body temperature was maintained at 37°C on a heating pad and a sterile “tips only” technique was utilized for all surgeries. A 10-µl Hamilton syringe connected to an UltraMicroPump and SYS-Mico4 controller (WPI, Sarasota, FL) was used to deliver 0.7 µl of AAV5-hSyn-hChR2(H134R)-eYFP (University of North Carolina vector core), over 5 min, into the medial VTA (AP: -5.4; ML: ±2.0; DV -8.2; 10° angle). After surgery, incisions were closed with absorbable sutures, and rats received an injection of the nonsteroidal anti-inflammatory meloxicam (1 mg/kg, s.c.) before being returned to their home cage where they were monitored for the next 3 days. Animals were euthanized for slice recordings 6-8 weeks following virus infusion.

#### Self-administration and LHb cannulae implantation

Surgical anesthesia was achieved with equithesin (1% pentobarbital, 2% magnesium sulfate, 4% chloral hydrate, 42% propyleneglycol, 11% ethanol, 3 ml/kg, i.p.), diluted 1:3 to 33% in saline immediately before injection to minimize peritonitis, and anesthetic depth was assessed continuously throughout the procedure. Rats were then stereotaxically implanted with bilateral guide cannulae (C315, Plastics One, Roanoke VA) aimed 1 mm dorsal to the LHb (coordinates: AP: -3.8, L: ±0.6, V: -3.6 mm relative to bregma). Rats for the Go/NoGo intravenous self-administration (IVSA) study were also implanted with a polyurethane catheter (Instech, PA) that was inserted 3.5 cm into the right jugular vein. The catheter terminated in a Vascular Access ButtonTM (Instech, PA) which was subcutaneously mounted in the animal’s back. Rats received an injection of the nonsteroidal anti-inflammatory meloxicam (1 mg/kg, s.c.) following surgery. Per ACUC policy, rats were monitored daily for signs of adynamic ileus; no such signs were observed in any subjects. All animals resumed normal feeding behavior and demonstrated weight gain during a one-week recovery phase prior to training in the Go/NoGo task.

### Go/NoGo Cocaine IVSA Task

Operant training took place as described in detail previously (Zapata et al., 2017). Standard rat operant chambers (Med-Associates; St Albans, VT) were used, and rats were trained to self-administer cocaine (0.75 mg/kg /infusion) for 12 sessions (2 hours or 40 infusions per session) on a schedule in which each lever press on the active lever was followed by an injection of cocaine (fixed ratio 1; FR1 schedule). Following this, rats were trained on a Go/NoGo task consisting of 2 hours sessions of 6 × 20 min alternating intervals of cocaine availability (Go) and non–availability (NoGo), signaled by the house light (light on during cocaine availability). During Go periods, every fifth response (FR5) was immediately followed by a cocaine infusion. During NoGo periods lever responses did not trigger cocaine infusions. Training progressed until stable discrimination of the Go/NoGo periods were observed (3 consecutive sessions in which NoGo responses were less than 30% of total, 12-14 sessions). After reaching criterion, testing proceeded using a within subject design in which each rat received bilateral infusions of PBS, scopolamine (50 mM), pirenzepine (30 mM), or AFDX-116 (30 mM), dissolved in PBS to a 0.5 µL injection volume.

### Data analysis

Statistical analyses were conducted using the two-tailed Student’s t-test and one- or two-way repeated-measures ANOVA (Prism 9, GraphPad Software, La Jolla, CA). An alpha value of *p* < 0.05 was considered statistically significant. The Sidak or Dunnett’s post-hoc tests were performed to assess between group differences.

## Results

### Effects of carbachol on LHb neuron membrane currents

We previously found that the muscarinic agonist oxotremorine-M (Oxo-M) produced heterogenous effects on LHb neuron somatic excitability (Zapata et al., 2017). Here, we measured the effect of the non-selective cholinergic agonist carbachol (CCh,10 µM) on LHb neuron membrane currents under voltage clamp (holding potential, I_hold_ = -60 mV). Like our previous Oxo-M dats, CCh reversibly activated either outward (inhibitory) currents (n = 10/31 cells from 20 rats; 32.3 %, **Figure 1B1**) or inward (excitatory) currents (n = 6/31,19.4 % **Figure 1C1**) in neurons located in the mLHb. The remaining 15 cells (48.4%) showed no change in I_hold_ upon CCh application. CCh-activated outward currents were associated with a reduction in input resistance (**Figure 1B2**), whereas no change in input resistance was observed in cells displaying inward currents (**Figure 1C2**). The inward holding currents were reversed by application of the M1R antagonist pirenzepine (PZP,1 µM; n = 4 cells from 3 rats; one way RM-ANOVA, F _2,6_ = 17.07, p = 0.003; CCh, p = 0.0027 vs. baseline; CCh + PZP, p = 0.4876 vs. baseline, Dunnett’s post-hoc). Although we were unable to obtain a large enough sample of outward currents to perform a detailed pharmacological analysis, pirenzepine failed to reverse outward currents in 2 cells examined (not shown).

**Figure 1.**
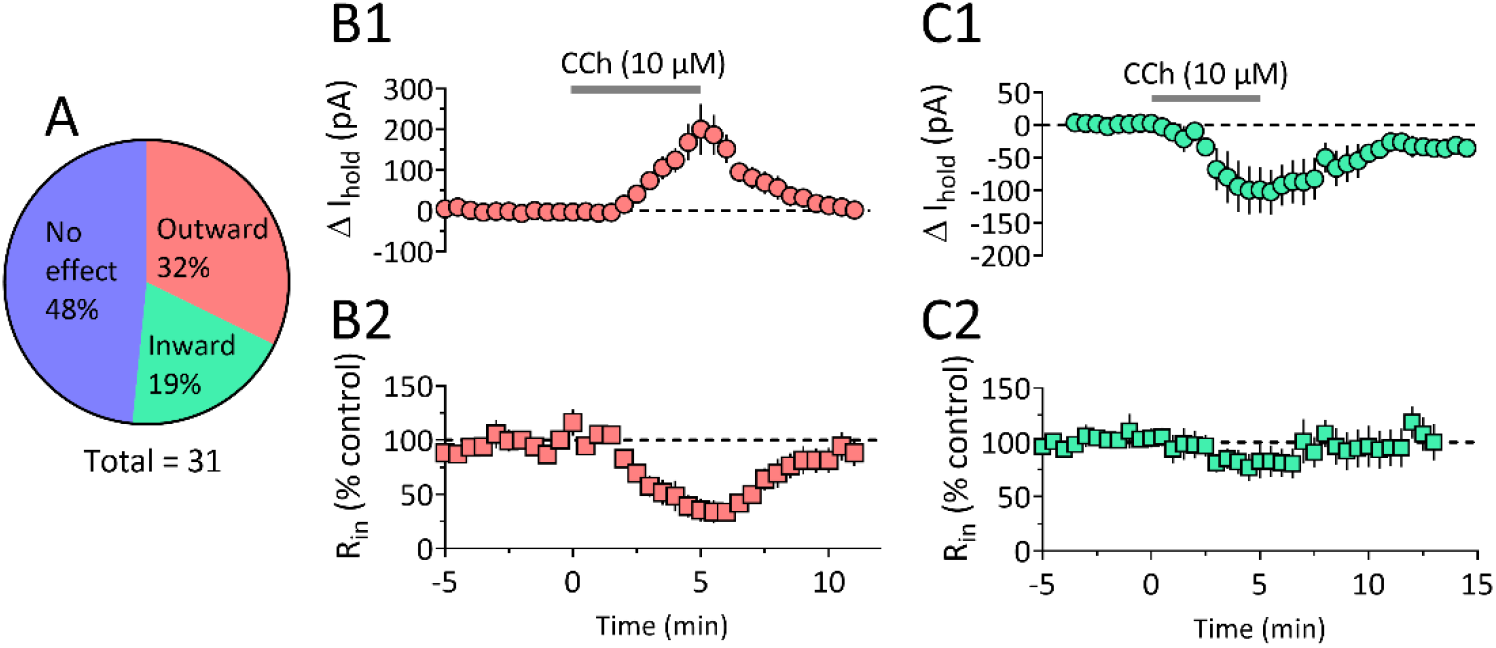
Somatic effects of carbachol on LHb neurons. (A) Proportion of cells demonstrating outward (n =10/31), inward (n = 6/31), or no change (n=15/31) in holding current (V_hold_ = -60mV) during bath application of carbachol (10 µM). (B1) Time course of mean outward holding current change (I_hold_), and (B2) associated decrease in cell input resistance during CCh application. (C1) Mean time course of inward holding current change, and (C2) no associated change in input resistance.

### Carbachol inhibits GABAergic synaptic input to LHb neurons via M2Rs

To determine whether mAChRs control synaptic integration in the LHb, we measured the effects of CCh on spontaneously occurring GABAergic IPSCs. Bath application of CCh (10 µM) significantly reduced the frequency of spontaneous IPSCs (sIPSCs) (**Fig. 2**; n = 13 cells from 4 rats, two-tailed paired t-test, t_12_= 4.6, p = 0.0006), but had no significant effect on the mean amplitudes of these events (two-tailed paired t-test, t_12_= 0.76, p = 0.46). This effect of CCh was not reversed by PZP (1 µM) (**Fig. 3C**; n = 7 cells from 4 rats; one-way RM-ANOVA, F _2, 12_ = 7.29, p = 0.0085; CCh, p = 0.005 vs. control; CCh + PZP, p = 0.04 vs control, Dunnett’s post-hoc). As high levels of expression of muscarinic M2 receptors (M2R) have been reported in the LHb (Wagner et al., 2016), we next evaluated the M2R antagonist AFDX-116 (Buckley et al., 1989; Auerbach and Segal, 1996) and found that it reversed the reduction in sIPSC frequency caused by CCh (**Fig. 3D**; 1 µM; one-way RM-ANOVA, F _2,12_ = 8.71, p = 0.005; AFDX-116, p = 0.15 vs control, Dunnett’s post-hoc). IPSCs evoked by local electrical stimulation (eIPSCs) of the brain slice were also significantly inhibited by CCh (**Fig. 4A, 4B**; one way RM-ANOVA, F _2, 18_ = 13.22, p = 0.003; p = 0.002 vs. control, Dunnett’s post-hoc), and this was also reversed by AFDX-116 (p = 0.24 vs. control, Dunnett’s post-hoc). The effects of CCh on eIPSCs did not differ by sex (males, n = 9, 44 ± 6% inhibition; females, n = 5, 31 ± 7% inhibition; two tailed t-test, t_12_ =1.4, p = 0.18). These data suggest that GABAergic inputs to LHb neurons are equally inhibited by presynaptic M2Rs in male and female rats.

**Figure 2.**
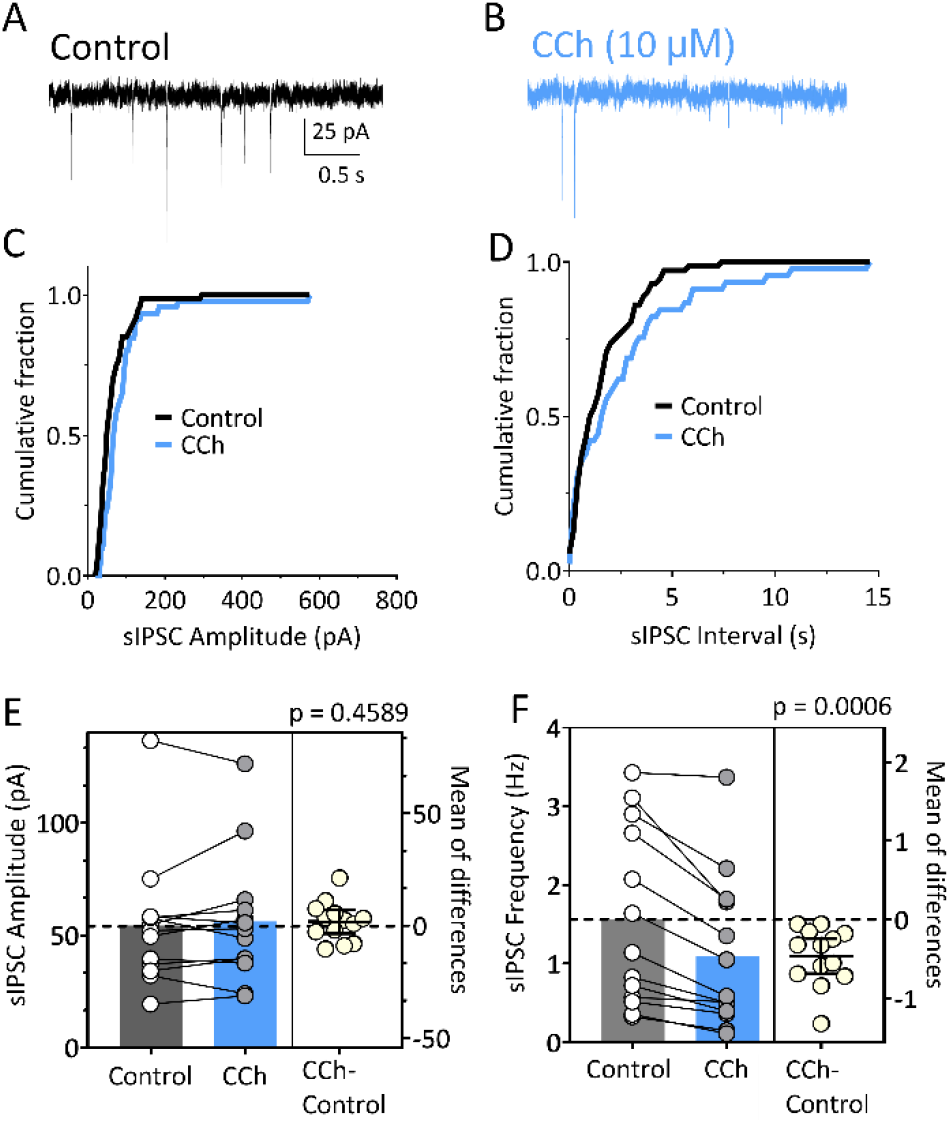
Carbachol inhibits spontaneous GABAergic IPSCs in LHb neurons. (A) Representative traces of sIPSCs under control conditions (V_hold_ = -70mV), and (B) during bath application of carbachol (CCh, 10 µM). Representative cumulative distribution histograms of the sIPSC amplitude (C) and inter sIPSC interval (D) from the same cell. (E) Estimation plot of sIPSC amplitude for all cells under control conditions and during CCh application. The difference for each cell between the control and CCh periods (CCh-control) is shown at right, with the mean difference ± 95% C.I. and statistical p value indicated. No significant difference in amplitude was observed (two-tailed paired t-test, t_12_ = 0.76, p = 0.46). (F) Estimation plot of sIPSC frequency for all LHb cells. A significant reduction in frequency was observed (two-tailed paired t-test, t_12_ = 4.6, p = 0.0006).

**Figure 3.**
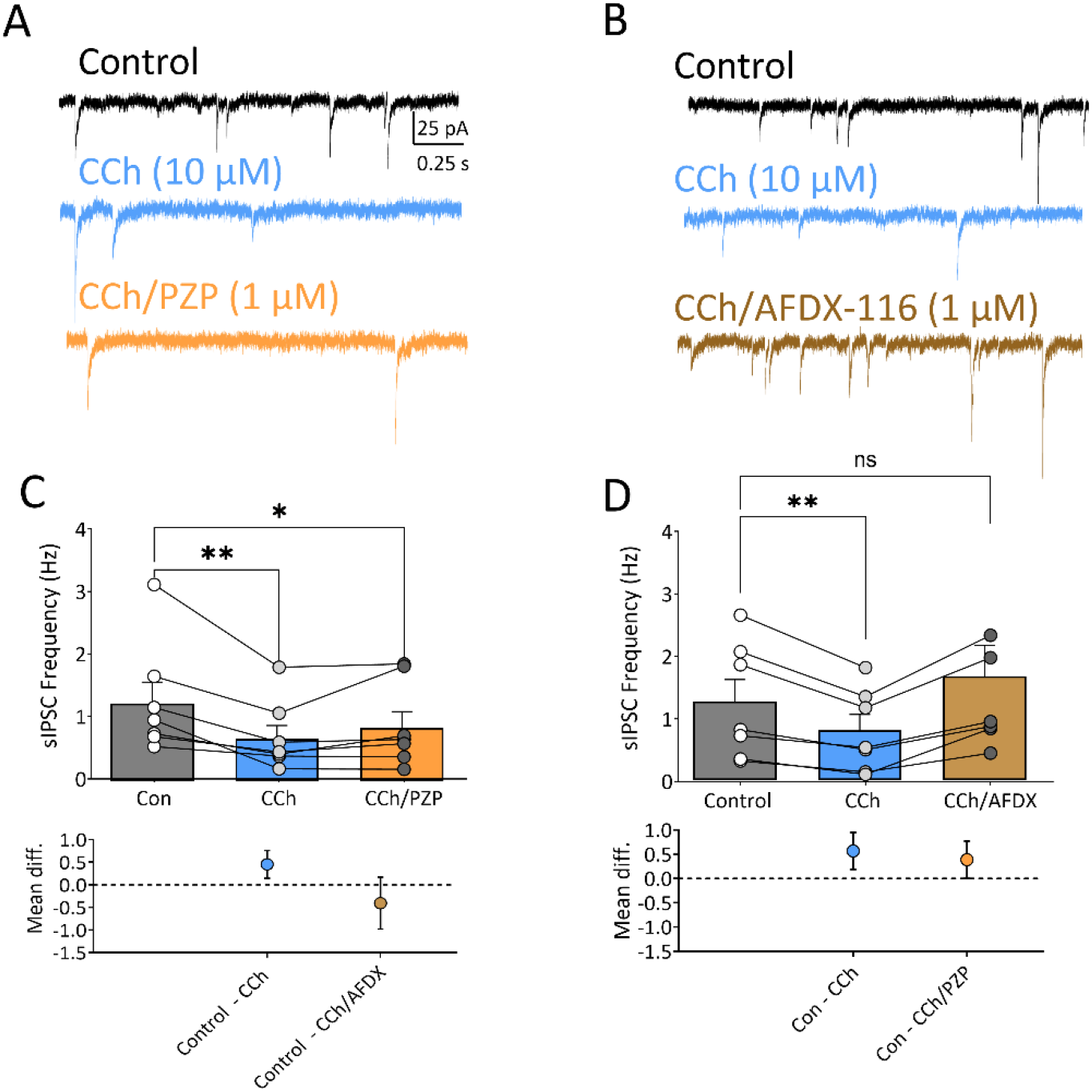
Carbachol inhibits spontaneous GABA release onto LHb neurons via activation of M2 muscarinic receptors. Representative sIPSC traces from individual LHb neurons showing the effect of carbachol (CCh, 10 µM), followed by co-application of (A) the M1 receptor antagonist, pirenzepine (PZP,1 µM) or (B) the M2 receptor antagonist AFDX-116 (1 µM). (C) PZP did not significantly reverse the effect of CCh on sIPSC frequency (one-way RM-ANOVA, F_2, 12_ = 7.29, p = 0.0085; CCh, ** p = 0.005 vs. control; CCh/PZP, * p = 0.04 vs. control, Dunnett’s post-hoc). (D) AFDX-116 significantly reversed the effect of CCh on sIPSC frequency (one-way RM-ANOVA, F_2,12_ = 8.71, p = 0.005; CCh, **p = 0.009 vs. control; CCh/AFDX, p = 0.15 vs. control Dunnett’s post-hoc). The graphs below the main panel show the mean difference (± 95% C.I.) for the post-hoc comparisons.

**Figure 4.**
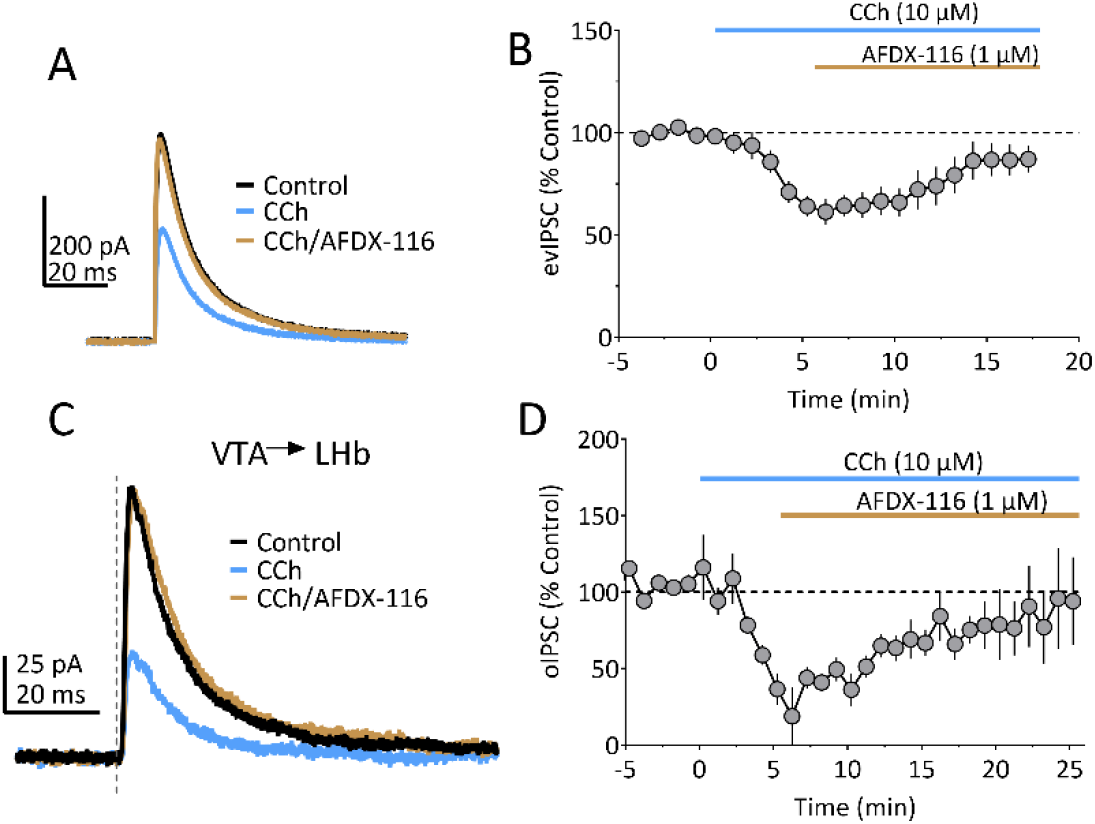
IPSCs evoked by local electrical stimulation and by light activation of ChR2 are inhibited by CCh acting at M2Rs in LHb. (A) Representative traces of electrically evoked IPSCs (V_hold_ = -45 mV, K-Glu based intracellular solution) showing inhibition during carbachol application (CCh, 10 µM) and reversal of this inhibition by co-application of the M2R antagonist AFDX-116 (1 µM). (B) Mean time course showing the effects of CCh and AFDX-116 on electrically activated IPSCs in LHb neurons (n =10 cells from 6 rats). Significant inhibition was observed following CCh (one way RM -ANOVA, F_2, 18_ = 13.22, p = 0.003; p = 0.002 vs. control, Dunnett’s post-hoc) and this was reversed by AFDX-116 (p =0.24 vs. control, Dunnett’s post-hoc). (C) Representative traces of light-evoked IPSCs recorded from a LHb neuron following infusion of synapsin-driven ChR2 into the VTA. The dashed line indicates the onset of a single, 5 ms duration light pulse (473 nm). (D) Mean time course of the effects of CCh and AFDX-116 on VTA to LHb light-evoked IPSCs (n = 5 cells from 2 rats). Significant inhibition was observed during CCh (one-way RM-ANOVA, F_2,8_ = 16.6, p = 0.001; p = 0.008 vs control, Dunnett’s post-hoc) and this was reversed by AFDX-116 (p = 0.09 vs. control, Dunnett’s post-hoc).

### Inhibition of GABAergic VTA inputs to LHb by M2Rs

As noted above, the LHb contains relatively few intrinsic sources of GABAergic inhibition (Brinschwitz et al., 2010; Aizawa et al., 2012; Wagner et al., 2017). Therefore, control of LHb neuron excitation likely arises from a diverse array of afferents, including those that arise from the VTA (Root et al., 2014b). As VTA neurons respond to both rewarding and aversive stimuli (Root et al., 2020), and express M2R mRNA (Vilaro et al., 1992), we hypothesized that this pathway would be sensitive to M2R-mediated synaptic control. Rats (n = 5) received VTA injections with AAV encoding for the channelrhodopsin (ChR2) protein, driven by a synapsin promoter (pAAV-hSyn-hChR2(H134R)-EYFP), and light-evoked optical IPSCs (oIPSCs) were recorded in LHb neurons 6-8 weeks later. The oIPSCs evoked by activation of ChR2 in VTA neuron axon terminals in the LHb were significantly inhibited by CCh (10 µM; **Fig. 4C, 4D**), and this was reversed by co-application of the M2R antagonist AFDX-116 (1µM; one-way RM-ANOVA, F_2,8_ = 16.6, p = 0.001; CCh, p = 0.008 vs control; CCh + AFDX, p = 0.09 vs. control, Dunnett’s post-hoc). These data suggest that VTA GABAergic inputs to LHb neurons can be inhibited by M2R activation.

### Carbachol inhibits glutamatergic synaptic input to LHb via M2Rs

We next examined whether synaptic glutamate input to LHb neurons is modulated by mAChRs by recording spontaneous EPSCs (sEPSCs). Unlike sIPSCs, CCh (10 µM) significantly decreased sEPSC amplitudes (**Fig. 5E**; p = 0.0002, t_31_ = 4.3, two-tailed paired t-test) and frequency (**Fig 5F**; p = 0.0002, t_31_ = 4.3, two-tailed paired t-test; n = 32 neurons from 18 animals), and this was independent of sex (two-way ANOVA; sex x treatment interaction, F_1,32_ = 0.3023, p = 0.58). Given that CCh hyperpolarizes some LHb neurons (**Fig. 1**), the reduction in sEPSC amplitude could reflect decreased somatic excitability of local glutamatergic LHb neurons that provide synaptic inputs to other LHb cells (Kim and Chang, 2005), as described for µ-opioids (Margolis and Fields, 2016). Therefore, to limit the influence of somatic excitability changes on glutamate release, we measured CCh effects on miniature EPSCs (mEPSCs) following blockade of action potentials by TTX (200 nM). In the presence of TTX, CCh reduced mEPSC frequency (n = 9 cells from 3 rats; control, 3.3 ± 0.39 Hz; TTX, 1.9 ± 0.23 Hz; **Fig. 5H;** two-tailed paired t-test, t_10_ = 5.36, p = 0.003) but had no effect mEPSC amplitude (control, 25.6 ± 4.1 pA; TTX, 24.9 ± 5.4 pA; **Fig. 5G**; two-tailed paired t-test, t_10_ = 0.51, p = 0.62). These results suggest that CCh can reduce synaptic glutamate release onto LHb neurons via changes in somatic excitability as well as by direct inhibition via effects at axon terminals.

**Figure 5.**
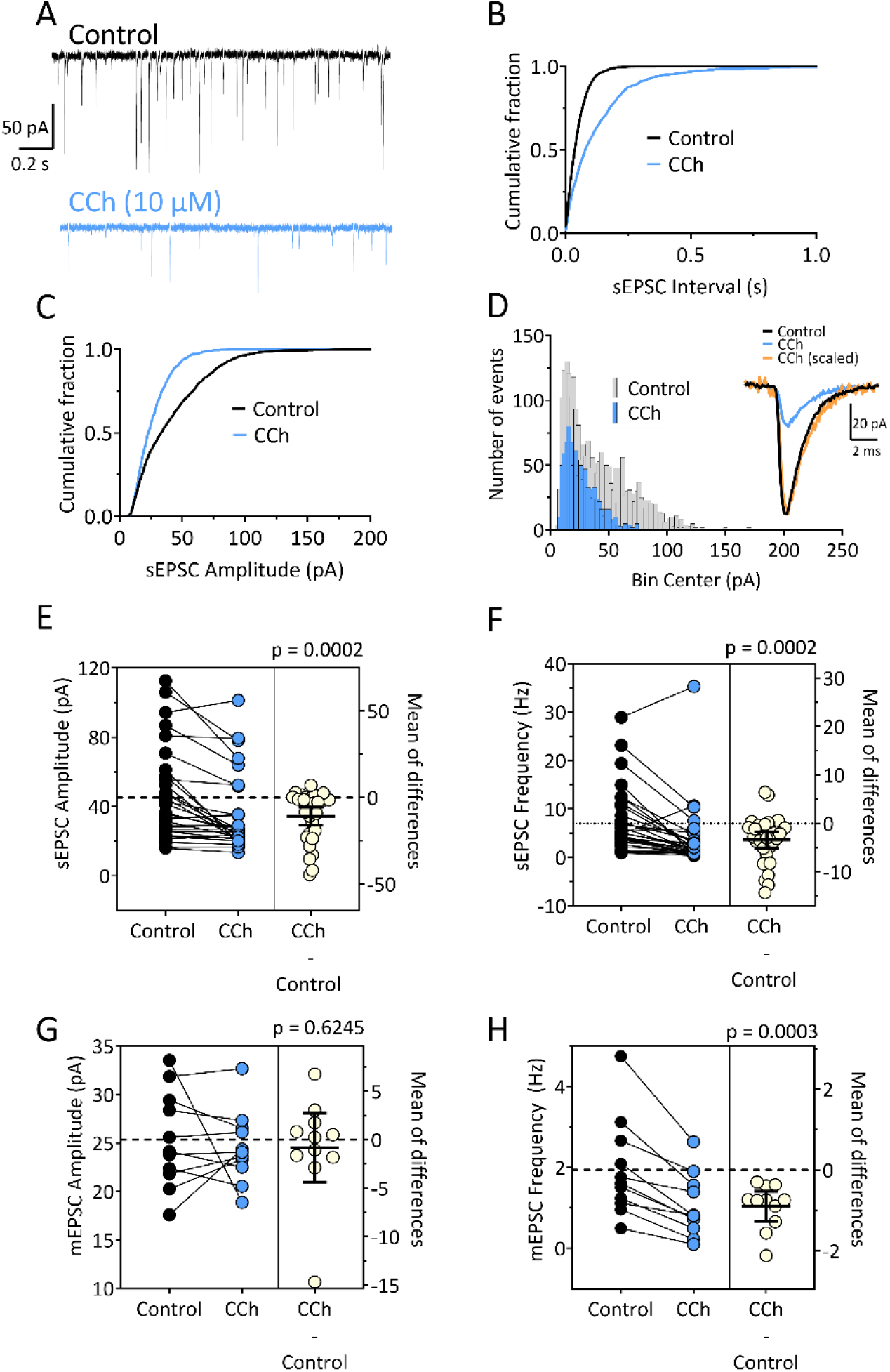
Carbachol inhibits glutamate release onto LHb neurons. (A) Representative traces of spontaneously occurring EPSCs (sEPSCs) under control conditions, and during bath application of CCh (10 µM). (B) Representative cumulative distribution histograms of inter-sEPSC intervals and amplitude (C) from the same cell demonstrate a reduction in the frequency and amplitude, respectively of the sEPSCs upon CCh application. (D) Non-cumulative amplitude histogram from the same cell demonstrates that the change in sEPSC amplitude results from a reduction in the number sEPSCs across the full range of amplitudes. The inset shows averaged sEPSCs in the absence and presence of CCh, as well as the waveform obtained in the presence of CCh scaled to the same size as the control waveform. This shows that CCh did not change in the rise or decay kinetics of the EPSCs. (E) Estimation plot of sEPSC amplitudes for every LHb neuron used in this analysis under control conditions and during CCh application. The right portion of the figure shows the difference in sEPSC amplitude between the control and CCh periods for each cell (CCh−control), and the mean difference ± 95% C.I. CCh caused a significant reduction in sEPSC amplitude (two-tailed paired t-test, t_31_ = 4.3, p = 0.0002). (F) Estimation plot of sEPSC frequency for each LHb neuron used in this analysis. A significant reduction in sEPSC frequency was observed following carbachol application (two -tailed paired t-test, p= 0.0002, t_31_ = 4.3, p = 0.0002). (G) The amplitude of mEPSCs, obtained in TTX (200 nM) was not significantly altered by CCh (10 µM) (two-tailed paired t-test, t_10_ = 0.51, p = 0.62). (H) mEPSC frequency measured in TTX was significantly reduced by CCh (two-tailed paired t-test, t_10_ = 5.36, p = 0.003).

Whereas the M1R antagonist PZP (1µM) did not affect the inhibition of sEPSC frequency by CCh (**Fig. 6A-6D**; one-way RM-ANOVA, F_2, 12_ = 6.396, p = 0.01; ** p < 0.01, Dunnett’s post-hoc), the M2R antagonist AFDX-116 (1µM) reversed the CCh-mediated inhibition of sEPSC amplitude and frequency (**Fig. 6E-6H**; one-way RM-ANOVA, F_2,14_ = 4.48, p=0.03; CCh, p = 0.02 vs. control; CCh + AFDX, p =0.6 vs. control, Dunnett’s post-hoc; one-way RM-ANOVA, F_2,14_ = 8.23, p=0.005; CCh, p = 0.003 vs. control; CCh +AFDX, p = 0.15 vs. control, Dunnett’s post-hoc). Thus, M2Rs, and not M1Rs, inhibit synaptic glutamate release onto LHb neurons.

**Figure 6.**
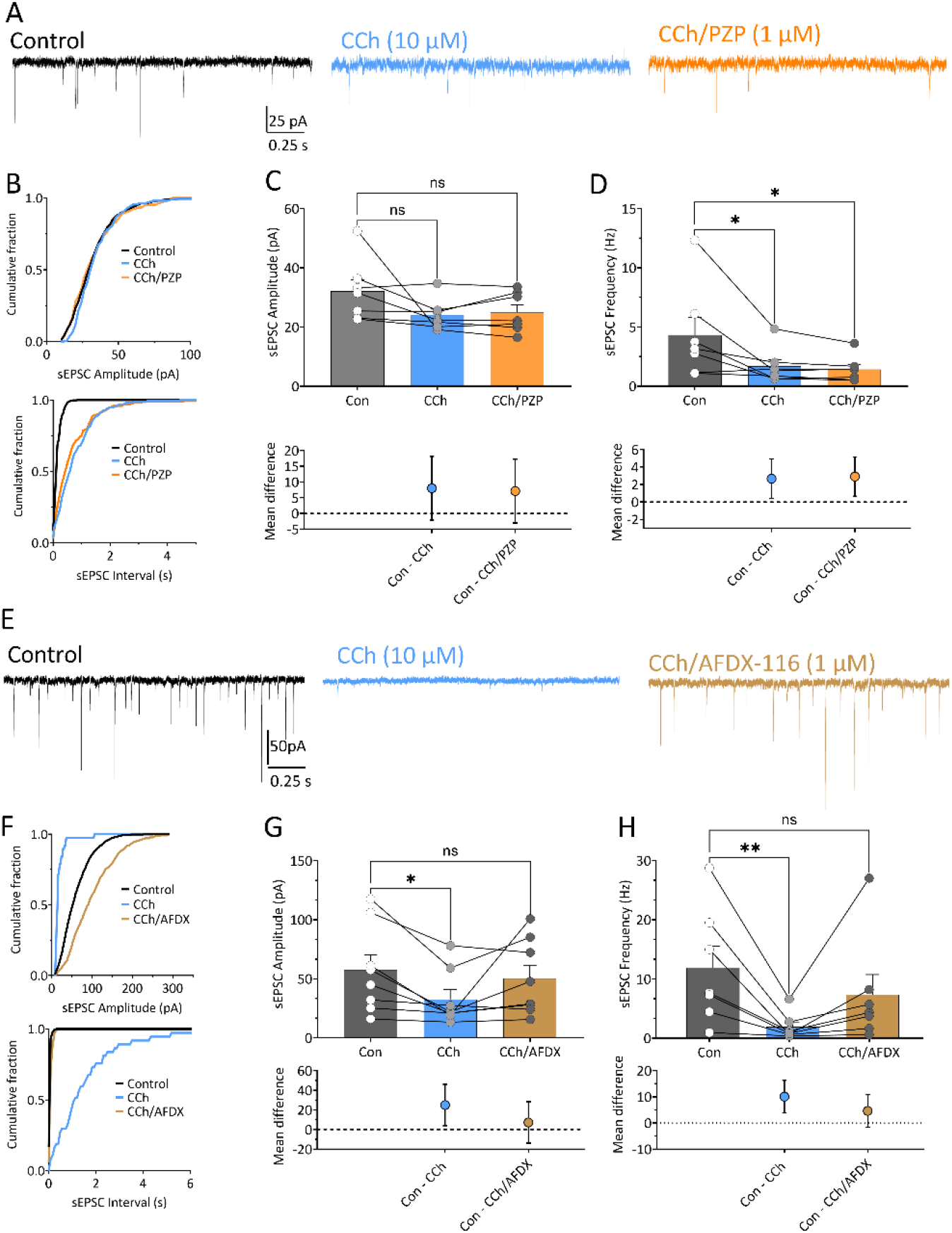
CCh inhibits glutamate release via M2 and not M1 mAChRs. (A) Representative sEPSC traces obtained under control conditions and during successive application of CCh (10 µM) and the M1R antagonist pirenzepine (PZP, 1 µM). (B) Cumulative distributions from the same cell for sEPSC amplitude (upper) and sEPSC interval (lower). In this cell, CCh did not alter the amplitude but clearly shifted the sEPSC interval. (C) sEPSC mean amplitudes are shown along with the mean differences in treatment vs. control (mean ± 95% C.I.). Although individual cells demonstrated reductions in amplitude, the mean effect was not significant (one-way RM-ANOVA, F_2, 12_ = 2.351, p = 0.14). (D) sEPSC frequency was significantly reduced by CCh, but this was not reversed by pirenzepine (one-way RM-ANOVA, F_2, 12_ = 6.396, p = 0.01; ** p<0.01, Dunnett’s post-hoc vs. control). (E) Representative sEPSC traces obtained under control conditions and during successive application of CCh and the M2R antagonist AFDX-116 (AFDX, 1 µM). (F) Cumulative distributions from the same cell for sEPSC amplitude (upper) and sEPSC interval (lower). Note the reversal of the effect following AFDX application. (G) Individual mean sEPSC amplitudes for each cell is shown. The mean amplitude was significantly reduced (one -way RM-ANOVA, F_2,14_ = 4.48, p=0.03; *p = 0.02 vs. control, Dunnett’s post-hoc) and this was reversed by AFDX (p = 0.6, Dunnett’s post-hoc). The mean difference (from control) and 95% C.I. for the treatment conditions are shown below. (E) sEPSC frequency was significantly reduced by CCh (one way RM-ANOVA, F_2,14_ = 8.23, p=0.005; **p = 0.003 vs. control, Dunnett’s post-hoc) and this was reversed by AFDX (p = 0.15 vs control, Dunnett’s post-hoc).

### M2Rs alter the excitatory-inhibitory balance of the LHb

The preceding data indicate that both excitatory and inhibitory synaptic inputs to LHb neurons can be inhibited by M2Rs. As an altered excitation-inhibition (E-I) balance in the LHb has been associated with a net change in neuronal activity and with dysfunctional reward valence attribution (Shabel et al., 2014; Meye et al., 2016; Mori et al., 2019), we next determined what the effect of M2R activation is on this E-I balance in the LHb (Antoine et al., 2019). CCh significantly reduced the E-I ratio measured using electrically-evoked EPSCs and IPSCs in the same LHb neurons (75 ± 10% of baseline; **Fig. 7**; n = 15 neurons from 7 animals; one sample t-test, t_14_ = 2.578, p =0.02). This reduction in E-I implies that, although both synaptic inhibition and excitation are both inhibited by CCh, excitatory glutamatergic inputs to LHb are reduced to a greater extent than inhibitory inputs. This change in the E-I ratio was also prevented by the M2 antagonist, AFDX-116 (1 µM; 13 neurons from 6 animals; 101 ± 6% of control, one sample t-test, t_12_ = 0.1098, p = 0.9144), and the effect of CCh on the reduction in E-I ratios in control versus AFDX-116-treated slices was significantly different (**Fig. 7B**; two tailed unpaired t-test, t_26_ = 2.16, p = 0.04). Thus, whereas both excitatory and inhibitory inputs to LHb neurons are sensitive to cholinergic inhibition, M2R activation inhibits excitatory inputs to a larger degree, thereby resulting in a net increase in the synaptic inhibition of LHb neurons.

**Figure 7.**
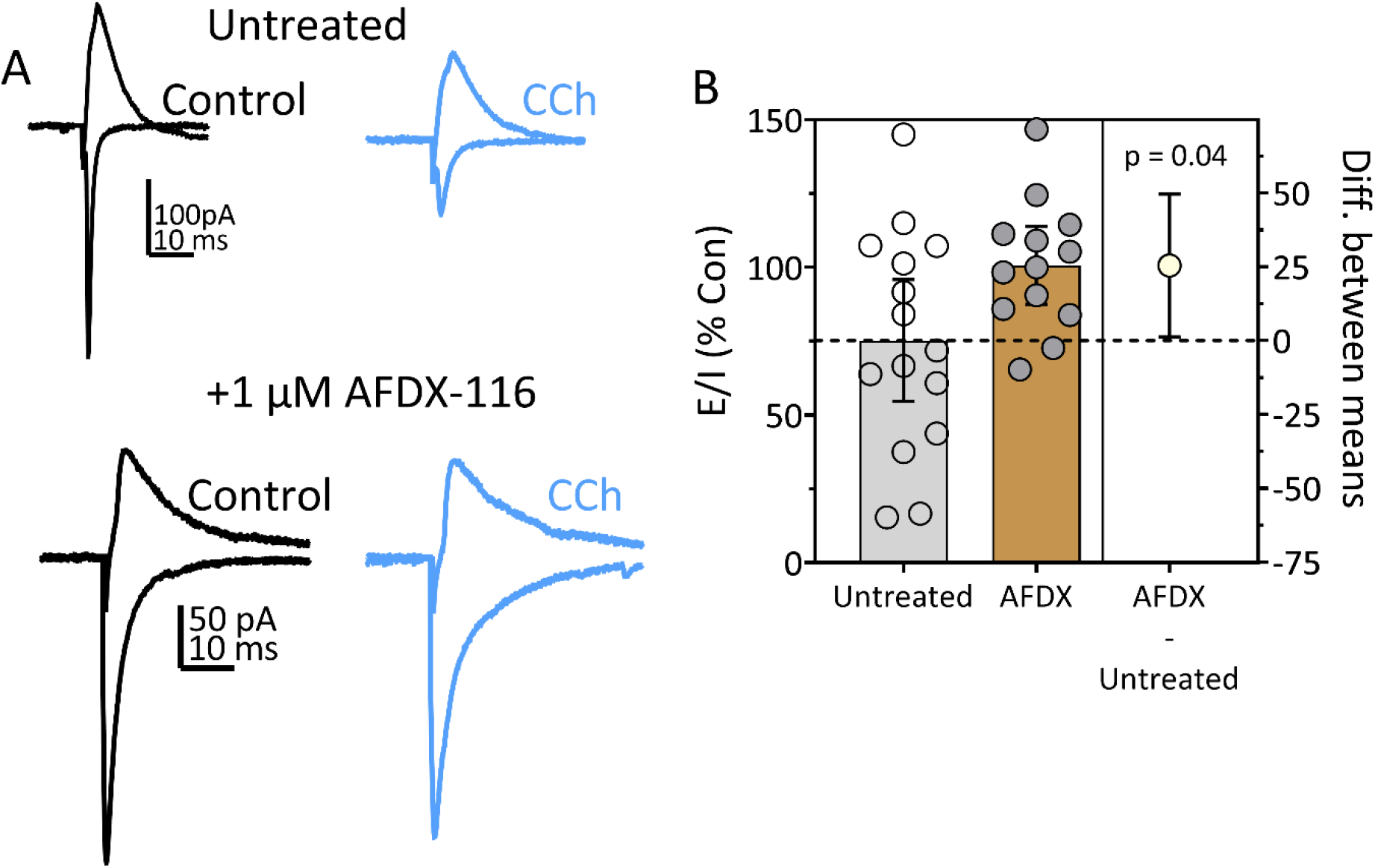
CCh reduces the excitatory/inhibitory (E/I) balance in LHb neurons via activation of M2Rs. (A) Traces from a single neuron before (control) and during CCh (10 µM) application. Electrically-evoked EPSCs (inward currents) were obtained at -70mV, and IPSCs (outward currents) were obtained at 0 mV in the same cell at the same electrical stimulus intensity. During CCh application, the E/I ratio, calculated as E/E+I using the total area of each response, was reduced. The lower panel shows IPSC and EPSC currents obtained from a different LHb neuron from a slice pretreated with the M2R antagonist AFDX-116 (1 µM). CCh had no effect on the responses or the E/I ratio. (B) Estimation-plot summary of the effect of CCh on the E/I ratio for all cells. The mean (± 95% C.I.) percentage reduction from the control E/I ratio produced is shown by the bar graph. The difference between the untreated and AFDX-treated slices was statistically significant (two tailed unpaired t-test, t_26_ = 2.16, p = 0.04).

### Intra-LHb blockade of M2Rs promotes impulsive cocaine -seeking

We previously reported that block of LHb mAChRs by the non-selective antagonist scopolamine impaired response inhibition for cocaine in rats trained on a Go/NoGo operant task (Zapata et al., 2017). This suggests that intact mAChR signaling is necessary for behavioral control in this task. Our present results demonstrate that M1Rs and M2Rs can alter LHb neuron membrane potential and that M2Rs alter synaptic integration in these cells. Therefore, we asked whether M1Rs or M2Rs were involved response inhibition in the Go/NoGo paradigm *in vivo*. Rats trained to self-administer cocaine and withhold operant responding during periods of signaled drug absence received bilateral infusions of vehicle, scopolamine (50 mM), PZP (30 mM), or AFDX-116 (30mM) prior to each test session. None of the intra-LHb infusions significantly altered responding during Go intervals when responses were rewarded with cocaine injections (**Fig. 8A;** RM-ANOVA, F_3,24_ = 0.6229, p = 0.6071). However, intra-LHb infusion of either scopolamine or AFDX-116 significantly increased responding during NoGo periods (expressed as the % of total responses) when lever presses were not followed by cocaine injections (**Fig 8B**; F_3,24_ = 7.983, p = 0.0060, RM-ANOVA; scopolamine, p = 0.00462 vs. Veh; AFDX, p = 0.0496 vs. Veh, Sidak’s post-hoc). In contrast to the effects of AFDX-116 on NoGo responding for cocaine, the M1R antagonist PZP did not significantly alter the response inhibition for cocaine during NoGo periods (**Fig. 8B**; p = 0.9197 vs Veh, Sidak’s post-hoc). Additionally, when the data were analyzed as the mean number of responses emitted during Go and NoGo periods we found that scopolamine and AFDX-116 specifically increased cocaine seeking responses during the NoGo intervals (**Fig 8C**; two-way RM-ANOVA, Go/NoGo vs scopolamine interaction, F_1,8_ = 8.264, p = 0.0207; **Fig 8D**; two way RM-ANOVA, Go/NoGo vs AFDX interaction, F_1,8 =_ 8.406, p = 0.0199). In contrast, PZP did not significantly affect the mean number of responses for cocaine during the Go or NoGo periods (**Fig 8E**; two-way RM-ANOVA, Go/NoGo vs PZP interaction, F_1,8_ = 0.3536, p = 0.5685). These data show M2Rs, and not M1Rs, are critical for response inhibition of cocaine seeking in the Go/NoGo task.

**Figure 8.**
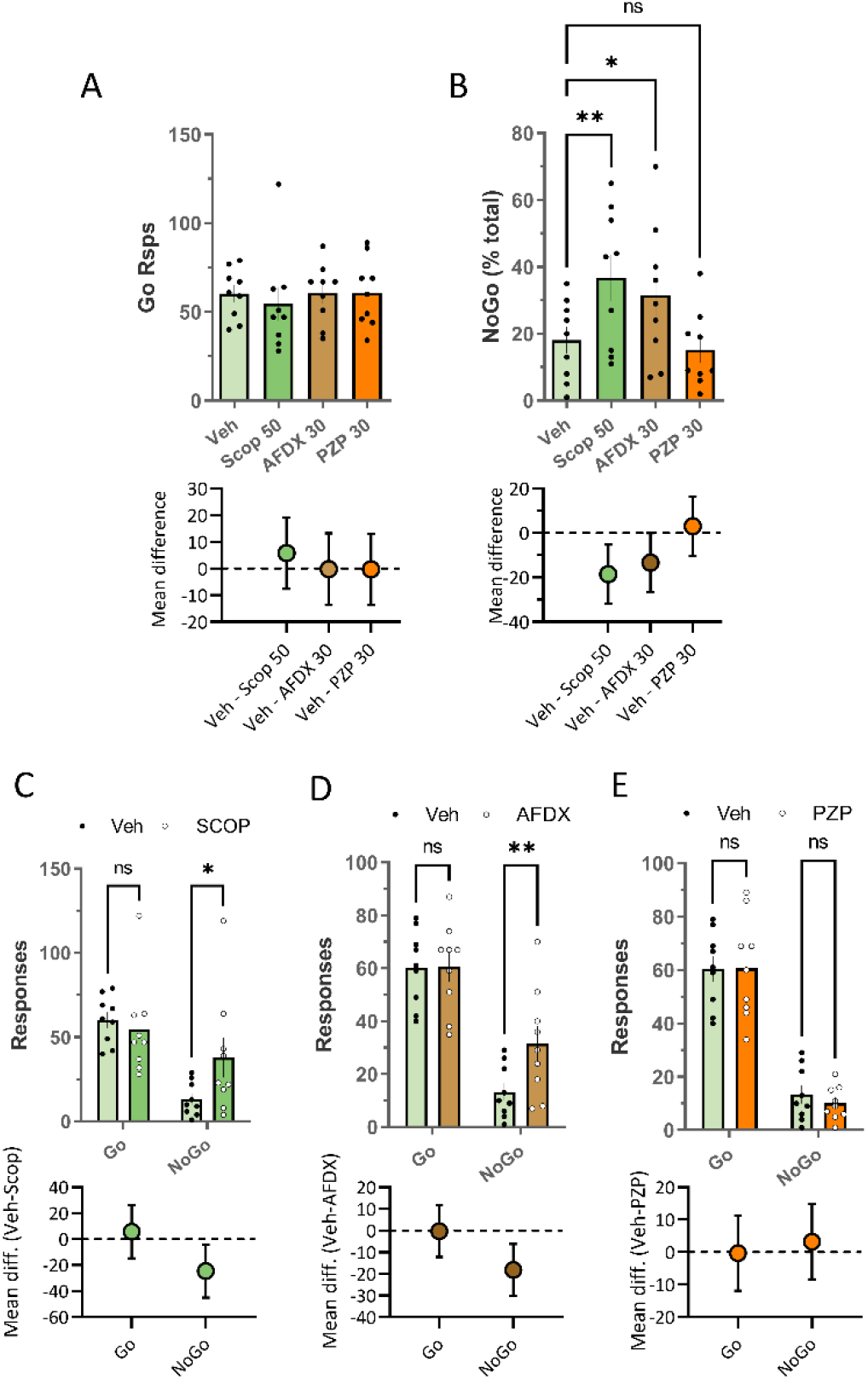
Blockade of LHb M2R mAChRs impairs inhibition of responses for cocaine in a Go/NoGo cocaine self-administration task. (A) Responses during signaled cocaine availability (Go responses) are unaffected by scopolamine (Scop, 50 mM), AFDX-116 (AFDX, 30 mM), or pirenzepine (PZP, 30mM). RM-ANOVA, F_3,24_ = 0.6229, p = 0.6071. (B) NoGo responses, as a percentage of total responses, are significantly increased by Scop (50 mM) and AFDX (30 mM), relative to vehicle (Veh; F_3,24_ = 7.983, p=0.0060, RM ANOVA; scop, p = 0.00462 vs. Veh; AFDX, p = 0.0496 vs. Veh, Sidak’s post-hoc). PZP (30 mM) did not significantly alter responding (p = 0.9197 vs Veh, Sidak’s post-hoc). Responses during Go and NoGo periods are shown for (B) scopolamine, (C) AFDX, and (D) pirenzepine. Significant increases in responding during NoGo periods, relative to Veh infusions, are indicated (two way RM-ANOVA, *p = 0.0219; **p = 0.0064, Sidak’s post-hoc). For all graphs, post-hoc differences (mean ± 95% C.I.) are shown in the lower panel.

## Discussion

Here we describe strong modulation of neurons in the parvocellular and central subnuclei of the mLHb by mAChRs and establish that M2Rs and the central cholinergic system are critical for operant response inhibition. We show that synaptic GABA and glutamate inputs to LHb neurons are inhibited by M2Rs and using optogenetics we demonstrate that a VTA to LHb projection represents at least one M2R-sensitive pathway. Whereas M2Rs inhibited both glutamate and GABA release onto LHb neurons, the net effect of M2R activation was to bias the E-I ratio toward enhanced inhibition. We predict this would reduce LHb excitatory output to downstream targets.

In addition to the altered synaptic integration by M2Rs, we also find effects of mAChRs on LHb neuron membrane currents that support the idea that M1Rs depolarize LHb neurons, whereas M2Rs seem to predominantly couple to inhibitory membrane currents. Although our behavioral study indicated that M2Rs are responsible for cholinergic control of response inhibition, we do not know whether this effect requires participation of presynaptic, postsynaptic, or both groups of M2Rs in the control of LHb output. Together, our data suggest that LHb mAChR signaling is critical for behavioral control involving response inhibition in drug seeking; whether these receptors control response inhibition in other behavioral contexts remains to be determined.

In a previous study, we reported that the mAChR agonist oxo-M depolarized approximately 50% of LHb neurons and hyperpolarized another 10% of these cells (Zapata et al., 2017). Here, CCh was used as an agonist because it exhibits more rapid pharmacokinetic properties in brain slices. However, we also found that, in contrast to oxo-M, CCh initiated outward currents (hyperpolarizations) in a larger proportion of LHb neurons (32.3%), and had no postsynaptic effect on a larger number of cells (48%). These differences likely result from the use of different agonists with distinct pharmacokinetic properties. However, the CCh-activated outward currents were associated with reduced input resistance, which is consistent with the opening of G-protein coupled inward rectifying potassium (GIRK) channels that are heterogeneously expressed in the LHb (Geisler et al., 2003; Zhang et al., 2016). As lower levels of these channels are found in the parvocellular and central subnuclei of the LHb (Geisler et al., 2003; Zhang et al., 2016), where most of the present recordings were performed, this may explain some of the heterogeneity in the response to mAChR activation we observe. Whereas a lower incidence of these postsynaptic effects precluded their more thorough pharmacological characterization, we found that CCh-activated depolarizing inward currents were reversed by M1R antagonism. Thus, despite the differences in the effects of CCh versus Oxo-M, mAChR activation can inhibit or excite different populations of LHb neurons.

In contrast to the heterogeneity in somatic response to CCh, more consistent synaptic effects in LHb neurons were observed. As sIPSC frequency was reduced by CCh in the absence of altered sIPSC amplitudes, the data suggest that mAChRs on inhibitory axon terminals reduced the probability of GABA release onto LHb neurons. Moreover, as this was reversed by AFDX-116 (Hulme et al., 1990), and not by pirenzepine, activation of M2Rs and not M1Rs inhibits GABA release in the LHb. IPSCs that were evoked by electrical stimulation of GABAergic axons arising from undefined inputs were also inhibited by M2Rs, as were those evoked by light-activation of ChR2 expressed in VTA afferents to LHb.

As the LHb lacks a major source of intrinsic GABA neurons (Brinschwitz et al., 2010; Aizawa et al., 2012; Wagner et al., 2017), the input from the VTA represents a major inhibitory afferent thought to strongly control LHb neuron output (Shabel et al., 2014; Barker et al., 2017; Faget et al., 2018). High levels of M2R expression have been detected in the LHb and M2R mRNA has been identified in projections to LHb originating in the VTA (Vilaro et al., 1992; Wagner et al., 2016). As VTA neurons projecting to the mLHb are known to co-release GABA and glutamate (Root et al., 2014b), and to encode both reward and aversion (Root et al., 2020), it will be important for future studies to evaluate the possibility that M2Rs can also inhibit glutamate release from these afferents and to determine how presynaptic M2Rs regulate behavior.

Glutamatergic synaptic inputs to LHb of unknown origin were also inhibited by M2Rs, as demonstrated by a significant reduction of sEPSC frequency and amplitude by CCh, and antagonism of this effect by AFDX-116. Although LHb neurons do not demonstrate extensive axonal collaterals (Weiss and Veh, 2011), there is evidence for some local connectivity among these glutamatergic cells (Kim and Chang, 2005). As we also found that mAChRs inhibit some LHb neurons, we reasoned that the reduction in sEPSC amplitude by CCh could reflect this somatic hyperpolarization, as described for µ-opioid receptors in the LHb (Margolis and Fields, 2016). Consistent with this, TTX pretreatment blocked the effect of CCh on sEPSC amplitude, but not sEPSC frequency, suggesting that CCh caused a reduction in larger action-potential dependent sEPSCs in the absence of TTX. The decrease in sEPSC frequency and amplitude by CCh was blocked by AFDX-116, suggesting that M2Rs can influence glutamate release by actions on both somatic excitability and, more directly, on the axon terminal glutamate release process. However, given that at least two major pathways to the LHb co-release GABA and glutamate (Root et al., 2014b; Shabel et al., 2014), we cannot exclude the possibility that some of the M2R-mediated inhibition of glutamate and GABA release occurs on these axons. Additional studies will be required to evaluate the extent to which M2Rs influence co-release from these inputs as well as focus on the influence of M2Rs on modulating somatic excitability within the LHb.

As M2Rs inhibited both glutamatergic and GABAergic synaptic transmission in the LHb, one might ask what the net of effect of this dual modulation might be? To investigate this, we examined CCh effects on E-I balance by measuring electrically evoked EPSCs and IPSCs in the same neurons. We found that CCh significantly decreased the E-I balance, resulting in a net shift toward increased inhibition of LHb neurons, and this was blocked by AFDX-116. Within the context of the indirect influence the LHb has on midbrain DA neurons via its output to the RMTg (Jhou et al., 2009), our data would predict that global activation of LHb M2Rs would dampen LHb excitation of RMTg to reduce inhibition of DA neurons and increase their excitability. However, this *in vitro* measurement of the effect of CCh on E-I balance occurs without consideration of the relative activity and contributions of these pathways in the behaving organism. Therefore, *in vitro* E-I balance can only provide a rough estimate of the contribution of M2Rs to LHb output and additional *in vivo* studies are required to more fully understand the contributions of mAChR-regulated neurotransmitter release to behavior. In this regard, our present results show that inappropriate NoGo cocaine seeking behavior increases during LHb M2R blockade. However, our prior study also showed that blockade of LHb glutamatergic receptors had no effect on this behavior (Zapata et al., 2017). These results therefore suggest that M2Rs on inhibitory afferents may be more important for response inhibition, and to drug seeking. Together the results of the present study, and our previous work, suggest that cholinergic signaling may represent an important target to influence LHb function and to modify its contribution to drug seeking, SUDs, and other psychiatric illnesses.

## Acknowledgements

This research was supported by the National Institutes of Health and the National Institute on Drug Abuse Intramural Research Program Grant, 1ZIADA000457 (C.R.L.).

